# Splice switching oligonucleotide mediated gene knockdown in B cells and plasma cells

**DOI:** 10.1101/2020.09.18.302984

**Authors:** Anne Marchalot, Jean-Marie Lambert, François Boyer, Justine Pollet, Jeanne Moreau, Jean Feuillard, Nathalie Faumont, Laurent Delpy

## Abstract

The need to identify new therapeutic approaches to the treatment of cancers of the B lymphoid lineage is crucial. Unlike CRISPR/Cas technology, antisense strategies result in transient modifications of gene expression and lack mutagenic effects at the DNA level. Here, we provide evidence for efficient knockdown of c-REL and RELA expression after treatment with splice switching antisense oligonucleotides (SSO) inducing exon skipping and reading frameshifts. We also developed a tool to facilitate the choice of exons for on purpose inhibition of mouse and human gene expression. Interestingly, treatments with morpholino SSO targeting the c-REL exon 2 donor splice site or RELA exon 5 acceptor splice site elicited very efficient knockdown in diffuse large B cell lymphoma (DLBCL) cell lines and antibody-secreting cells derived from primary human B cells. Consistent with the clinical relevance of c-REL activation in DLBCLs, treatment with c-REL SSO induced major alterations in NF-κB and TNF signalling pathways and strongly decreased cell viability. Altogether, SSO-mediated knockdown is a powerful approach to transiently inhibit the expression of given genes in B-lineage cells that should pave the way for cancer treatments, provided optimized ligand-conjugations for *in vivo* delivery.

**Graphical Abstract:** 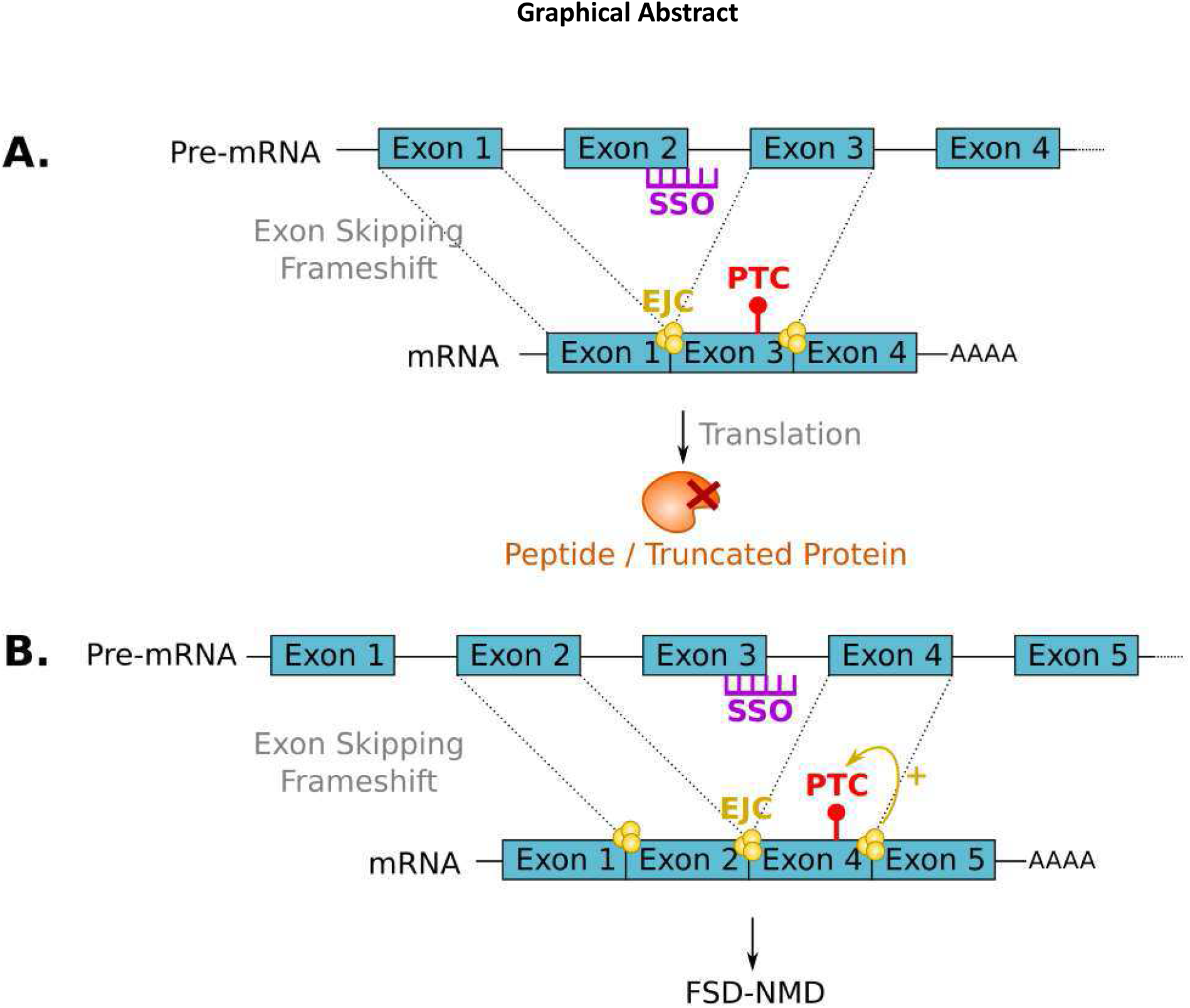

## INTRODUCTION

B cells and plasma cells are key actors in humoral immune responses by producing antibodies that provide long-term defense against pathogens such as viruses, bacteria or fungi. B cells are activated when extracellular antigens bind their B cell receptor (BCR) which induces rapid cell proliferation and differentiation into antibody secreting cells (ASC), either plasmablasts or plasma cells, generating sustainable antibody production (1). However, even if this mechanism is well described in literature, some questions remains concerning the B cell differentiation process in humans, notably how plasma cells home to their bone marrow niches (2), how high affinity clones are selected (3) and what mechanism lies under their extended longevity (4,5). Unravelling these questions could give new leads to treat antibody-mediated diseases such as allergy, autoimmunity or multiple myeloma and even benefit vaccine development. Studying plasma cell differentiation using human *in vitro* models rather than mouse models which differ significantly in this area (6) and where some ASC related diseases still cannot be fully reproduced (7–9) seems to be the best approach. However, gene knockdown in primary human plasma cells and B cells has always been a struggle, impeding research in non-cancer human B cells and plasma cell studies. Indeed, RNAi mediated gene knockdown in primary B cells is known to induce high mortality rates and poor penetrance during transfection, responsible for a lower knockdown efficiency compared to transformed B and plasma cells (10). Recently, it was demonstrated that CRISPR-Cas9 technology is efficient for gene knockout in expanded primary B cells and plasma cell differentiation models (11,12) but additional tools to realize a transient knockdown and avoid DNA modifications that could affect other genes (13,14) are still needed.

Antisense oligonucleotides (ASO) are short single chains of modified DNA that can be used to degrade mRNA by RNAse H1 recruitment or to modify pre-mRNA splicing among other mechanisms, depending of the oligonucleotide chemistry (for review see (15,16)). In addition to RNAse H1-dependant RNA degradation, splice-switching oligonucleotides (SSO) can also be designed to knockdown gene expression, as shown for STAT3 by Zammarchi and colleagues (17). These authors demonstrated that splice switching oligonucleotides (SSO) could be used to induce forced splicing-dependent nonsense-mediated decay (FSD-NMD), a mechanism based on SSO-mediated exon skipping to create a reading frameshift, the creation of a premature stop codon (PTC) and ultimately degradation of alternative mRNAs by NMD (17). While any splicing modulation producing a reading frameshift and the appearance of a PTC can be suitable for FSD-NMD, exon-skipping is the most frequent alternative splicing event in humans (18) and can be easily achieved with SSO hybridizing to either the donor splice site (5’SS), the acceptor splice site (3’SS) or exonic or intronic splicing enhancer (ESE or ISE) sequences (19). However, to efficiently knockdown gene expression, the delivery of sufficient SSO amounts into intracellular compartments remains challenging, as for RNAi technology. Over the last decades, numerous strategies have been developed to overcome this limitation and molecules such as peptide/ligand conjugates, nanoparticles or adeno-associated viruses (AAV) have been used to improve the delivery of antisense compounds (20). However, the development of such FSD-NMD strategies to modulate gene expression in B-lineage cells, including primary B or plasma cells, is still lacking.

During plasma cell differentiation, some key genes are differentially regulated including Bcl-6 and Blimp1 expressed in B cells and plasma cells respectively (21). Among those genes, NF-κB components such as c-Rel and RelA are respectively down- and up-regulated during plasma cell differentiation. Blimp1 is upregulated by RelA expression which in turn, downregulates c-Rel expression by binding its enhancer in ASC (22). Moreover, c-Rel and RelA expression is upregulated in many cancers, especially those of B cell lineage origin (23,24). Therefore, c-Rel and RelA could be interesting targets to cure B cell and plasma cell malignancies like diffuse large B cell lymphoma (DLBCL) or multiple myeloma (MM).

In this study, we describe a pipeline for the design of SSO to knockdown gene expression, either by inducing FSD-NMD or by shortening the open reading frame into an ineffective short polypeptide. Then, we evaluated this method in B cells using SSO targeting c-Rel and RelA transcripts, by comparing morpholino SSO transfection to passive vivo-morpholino administration and finally, we verified the efficacy of this strategy in a plasma cell differentiation model.

## MATERIAL & METHODS

### ASO design

REL and RELA gene sequences were collected from NCBI, GRCh38.p12 assembly, 25 base pairs upstream to 25 base pairs downstream from each splice site. SSOs targeting c-Rel exon 2 donor splice site (5′-ACAGAAGAAGGGTCTATTACCTGGA-3′), RelA SSO targeting the acceptor splice site of exon 5 (5′-GAAACTGAGCGCCCCCAGTCGTC-3′), and irrelevant Control ASO (5′-CCTCTTACCTCAGTTACAATTTATA-3′) were designed as morpholino or Vivo-morpholino and purchased from Gene Tools, LLC. 0.5mM ASO stock solutions were prepared with sterile nuclease-free water.

### Cell culture and ASO treatments

SUDHL4 and OCILY10 (DLBCL) cell lines were cultivated in RPMI1640 medium with Ultraglutamine (Lonza) and 10% or 20% Fetal Bovine Serum (FBS) (Dominique Dutscher) at 37°C with 5% CO2 and treated for 48 hours with 0.25 to 0.75 µM ASO for SUDHL4 and with 1.5 to 2 µM ASO for OCILY10. Human Peripheral Blood Mononuclear Cells (PBMC) were obtained by density gradient centrifugation (Lympholyte-H Celardane) of cytapheresis rings from healthy donors (EFS Bordeaux) and B cells were sorted with the B cell isolation kit II (Miltenyi Biotec). Sorted B cells were cultured for 4 days in RPMI 1640 with UltraGlutamine (Lonza) containing 10% FBS (Dominique Dutscher), 1 mM sodium pyruvate (Eurobio), 1% AANE (Eurobio) and 50 U/ml penicillin/ 50 µg/ml streptomycin (Gibco) with 1µg/mL CPG-ODN2006 (Miltenyi), 2,4µg/mL BCR Fab’2 IgA+G+M (Jackson Immunoresearch), 5ng/mL hIL-2 (R&D Systems) and 100ng/mL Mega-CD40 ligand (Enzo Life Sciences) then washed and stimulated in the same media with 5ng/mL hIL-2 (R&D Systems), 12ng/mL hIL-10 (R&D Systems) and 5ng/mL human IL-4 (Peprotech) for 3 more days. Human B cells were treated with 2µM ASO for 72 hours at day 1 following stimulation for plasmablast studies and at day 4 following stimulation for plasma cell studies.

### Flow Cytometry

Apoptosis in SUDHL4 cells was determined using AnnexinV antibody (BioLegend) and 7-AAD (BD Pharmingen) staining. Data were acquired on a Beckton Dickinson LSRII Fortessa cytometer and analyzed with FlowLogic software.

### Western Blot

Cells were lysed in radioimmunoprecipitation assay (RIPA) buffer (Thermo Scientific) supplemented with a protease and phosphatase inhibitor cocktail. Lysates were sonicated and protein concentrations were determined using Pierce™ BCA Protein Assay kit (Thermo Scientific). Proteins were denaturated at 95°C for 5 minutes before separation on SDS-PAGE TGX 12% Stain-Free FastCast Acrylamide (Bio-Rad Laboratories). Proteins were then electro-transferred onto Trans Blot Turbo polyvinylidene fluoride membranes (Bio-Rad Laboratories). Western blots were probed with rabbit anti-human c-Rel antibody (Cell Signaling), rabbit anti-human RelA antibody (Cell Signaling) or mouse anti-beta-actin antibody (Sigma). Detection was performed using an HRP-linked goat anti-rabbit or goat anti-mouse secondary antibody (Southern Biotech) and chemiluminescence detection (ECL Plus™, GE Healthcare) using ChemiDoc™ Touch Imaging System (Bio-Rad Laboratories).

### RT-PCR and qPCRs

Total RNA was prepared using Tri-reagent (Invitrogen) procedures. RT-PCR was carried out on 1µg DNase I (Invitrogen)-treated RNA using High-Capacity cDNA Reverse Transcription Kit (Applied Biosystems). PCRs were performed on cDNA samples equivalent to 10 to 20 ng of RNA per reaction using the Taq Core Kit (MP Biomedicals) and the following primer pairs: 1) forward 5’-GGCCTCCTGACTGACTGACT and reverse 5’-GTAGCCGTCTCTGCAGTCTTT for c-Rel and 2) forward 5’-GCGAGAGGAGCACAGATACC and reverse 5’-TCACTCGGCAGATCTTGAGC for RelA. PCR products were separated on 2% agarose TBE gels. Quantitative PCRs were performed on cDNA samples equivalent to 5-10 ng of RNA per reaction, using SYBR® Premix Ex Taq ™(Tli RNaseH Plus), ROX plus or Premix Ex Taq ™ (Probe qPCR), ROX plus (Takara) on a StepOnePlus Real-Time PCR system (Applied Biosystems). Transcripts were quantified according to the standard 2^-ΔΔCt^ method after normalization to GAPDH (Hs02758991_g1 ThermoFisher Scientific Probe). SYBR quantitative PCRs were performed for the quantification of c-Rel full length transcripts with forward 5’-GGCCTCCGGTGCGTATAA and reverse 5’-TGTTCGGTTGTTGTCTGTGC primers and for exon 2 skipped c-Rel transcripts with forward 5’-AGCCATGGCCTCCGATTATG and reverse 5’-AAGGTCTGCGTTCTTGTCCA primers.

### RNA sequencing

Messenger RNA-sequencing was performed on the Illumina NextSeq500 and analyzed with DESeq2 statistical analysis at Nice-Sophia-Antipolis Functional Genomics Platform. Differentially expressed gene were sorted when the adjusted p value was > 0.05 and the fold change was < 1.5 or > 1.5. Reads were aligned with STAR on the hg38 genome version during the primary analysis. REL read visualization was performed on IGV software (Broad Institute and the Regents of the University of California) and sashimi plots were obtained by the ggsashimi tool on R studio software with a cutoff of 5 junctions. Heatmaps were obtained with R studio software using ggplot tool on statistically and differentially expressed gene lists. Pathways enrichments were analyzed on statistically and differentially expressed gene lists with g:Profiler website (25).

### Transcriptomic analysis

The use of Python (26) package gffutils (27) was used to generate a database from the human genome annotation file (GTF format) of GENCODE 34 version (28). This database was used to search the number of exons among the coding sequence of each transcript. For figure 1B, coding sequences composed of less than 3 exons were excluded and only internal exons among these coding sequences, excluding first and last exons, were analyzed for the condition: number of nucleotides not divisible by 3 for each coding sequences. Results were written in a TSV file using Python libraries NumPy (29) and pandas (30) then used to generate pie charts with the Python graphic library Plotly (31).

**FIG 1.**
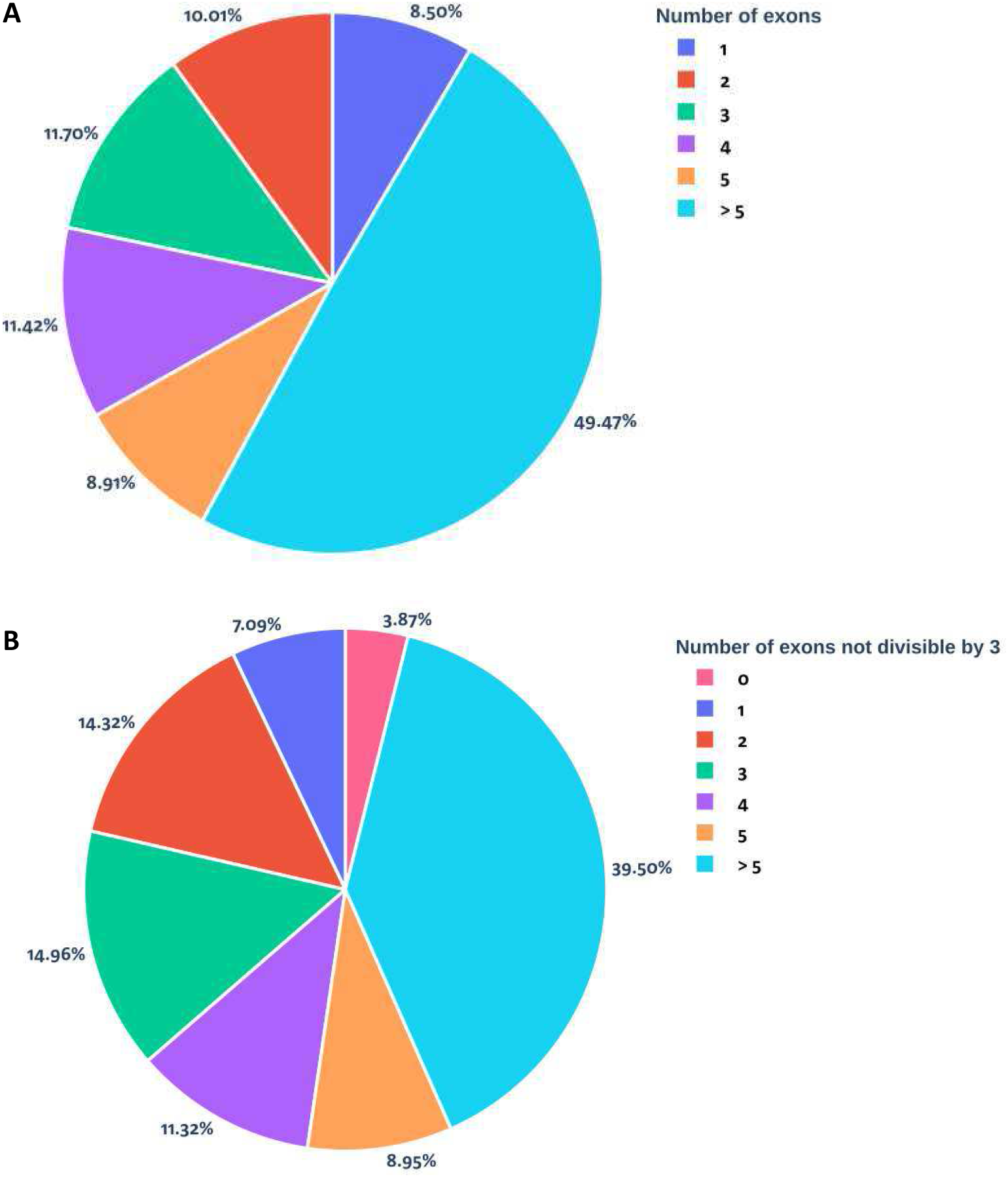
Human whole transcriptome analysis of coding sequences. **A**, Number of exons in the coding sequence **B**, Number of exons not divisible by 3 among coding sequences bearing at least 3 exons and excluding the first and last exons.

### Statistical analysis

Results were expressed as the mean ± standard error of the mean (SEM), and overall differences between variables were evaluated by an unpaired two-tailed Student’s t test using Prism GraphPad software (San Diego, CA).

## RESULTS

### Splice switching oligonucleotide selection workflow for exon skipping-mediated knockdown of gene expression

Splice switching oligonucleotide (SSO) mediated gene knockdown can be achieved with an exon-skipping strategy for the vast majority of human genes, containing ≥ 3 exons with a targeted internal one not divisible by 3 for a reading frameshift to occur. In addition, exon skipping must provoke the appearance of a PTC to drastically shorten the open reading frame and/or support NMD degradation. Transcriptomic analysis revealed that 81.5% of human coding sequences exhibited at least 3 exons (Figure 1A). Among these coding sequences bearing at least 3 exons, the number of coding sequence harbouring 1, 2, 3, 4, 5 and >5 internal exons not divisible by 3 were 7.09%, 14,32%, 14.96%, 11,32%, 8.95% and 39.50% respectively and only 3.84% were not eligible for this strategy (Figure 1B).

We developed an R shiny tool called Exon Skipper (https://cribl.shinyapps.io/ExonSkipper/) for rapid identification of putative exon candidates for any mouse or human transcripts. This tool also indicates the PTC position for each exon skipping event and provides predictive information with regards to NMD degradation and ORF shortening. For each selected exon candidate, we then examined its nucleotide sequence, together with its surrounding 500 nucleotide intronic and exonic sequences, to detect unwanted restoration of the reading frame using cryptic splice sites. This analysis can be performed by any splice site scoring tools available such as Human Splicing Finder (HSF) (https://hsf.genomnis.com/sequence) (32). The selection of 3’SS targeting SSOs is preferred when a high risk of cryptic 5’SS that restores the reading frame is found and conversely. In the absence of predictive correction of reading frames after alternative splicing involving cryptic splice sites, SSOs can be designed to hybridize to either 5’SS, 3’SS or ESE/ISE sequences; the latter can be identified using Skip-E (https://skip-e.geneticsandbioinformatics.eu), ESE finder (http://krainer01.cshl.edu/cgi-bin/tools/ESE3/esefinder.cgi) (33) or other tools. Each SSO sequence must be carefully chosen as there are optimal parameters for each oligonucleotide chemistry related to RNA binding affinity (for design tips see (19)). RNA binding parameters can be analysed through IDT OligoAnalyzer (https://eu.idtdna.com/calc/analyzer) and RNA availability through folding RNA simulators such as mfold (http://unafold.rna.albany.edu/?q=mfold) (34). Finally, NCBI BLAST (https://blast.ncbi.nlm.nih.gov/Blast.cgi) analysis of selected SSO sequences towards other transcripts from the same organism is required to exclude SSOs with a high risk of cross-hybridization to off-targets (Figure 2). This complete method referred to as “Exon skipper pipeline” was used for the design of SSOs targeting two members of the NF-kB pathway, c-Rel and RelA.

**FIG 2.**
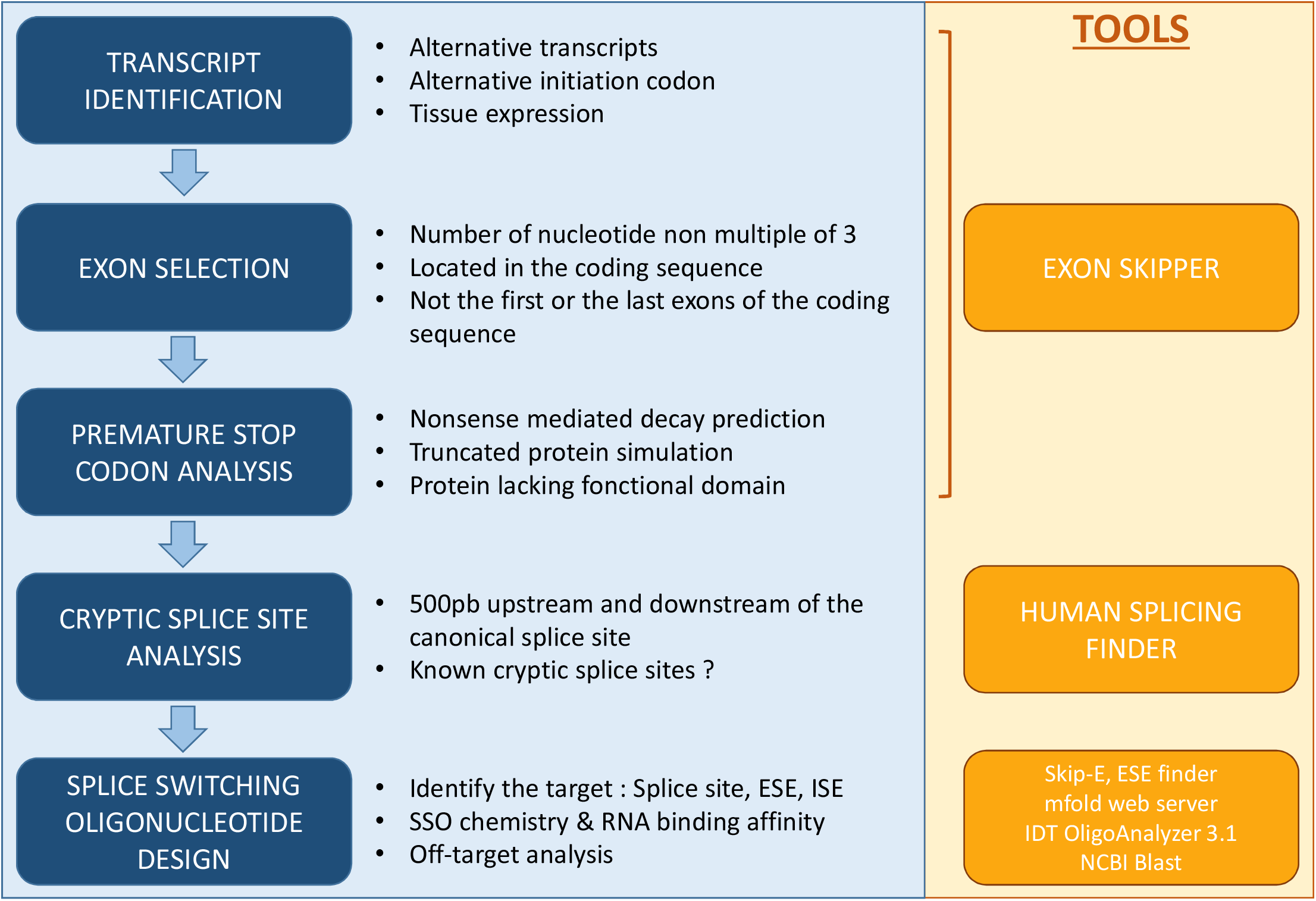
“Exon skipper pipeline” facilitates selection of splice switching oligonucleotides (SSO) to knockdown gene expression. Schematic representation of in-silico sequential steps performed to select exon targets and define the best SSO position for efficient inhibition of gene expression.

### SSO targeting of the c-Rel exon 2 donor splice site efficiently knocksdown c-Rel expression in B-lineage cells

An SSO targeting the exon 2 donor splice site of c-Rel was designed with the previously described method in order to convert c-Rel protein translation into an inactive 21 amino acid peptide, lacking all active domains compared to the full-length c-Rel isoform. Consistent with the NMD escape of PTC-containing mRNAs with short ORFs (35,36), alternative c-Rel mRNAs lacking exon 2 are likely to be poor NMD substrates as the PTC is close to the translation initiation site (35,36) (Figure 3A). First, SUDHL4 (DLBCL) cells were used to evaluate the efficacy of passive administration of vivo-morpholino SSO targeting c-Rel. Treatment with 0.25 to 0.75µM vivo-morpholino SSO for 48 hours was sufficient to induce a dose-dependent decrease in c-Rel expression (Figure 3B,C), with a 10-fold reduction of full-length c-Rel mRNA expression compared to irrelevant control treated cells (CTRL) (Figure 3B,C) and a nearly complete absence of c-Rel protein at a concentration of 0.5µM. As predicted, the c-Rel exon 2 skipped transcripts induced in SSO conditions were readily detectable by RT-PCR as they are predicted to be poorly degraded by NMD (Figure 3B). Comparatively, transfection of morpholino SSO was far less efficient than passive administration of vivo-morpholino SSO and did not achieve complete knockdown of c-Rel expression (Figure S1). We further verified these results by other experiments on SUDHL4 treated with 0.5µM SSO and confirmed an effective reduction of full-length c-Rel mRNA expression (Figure 3E,F). c-Rel SSO was then used in a plasma cell differentiation model described by Le Gallou et al (37,38) and 2µM efficiently provoked c-Rel protein knockdown in pre-plasmablasts where c-Rel is still highly expressed (Figure 3G,H) and also in plasma cells where its expression was reduced (Figure S2). Thus, we identified vivo-morpholino SSO targeting the c-Rel exon 2 donor splice site as simple and powerful tool to use for the inhibition of c-Rel expression in B and plasma cells.

**FIG 3.**
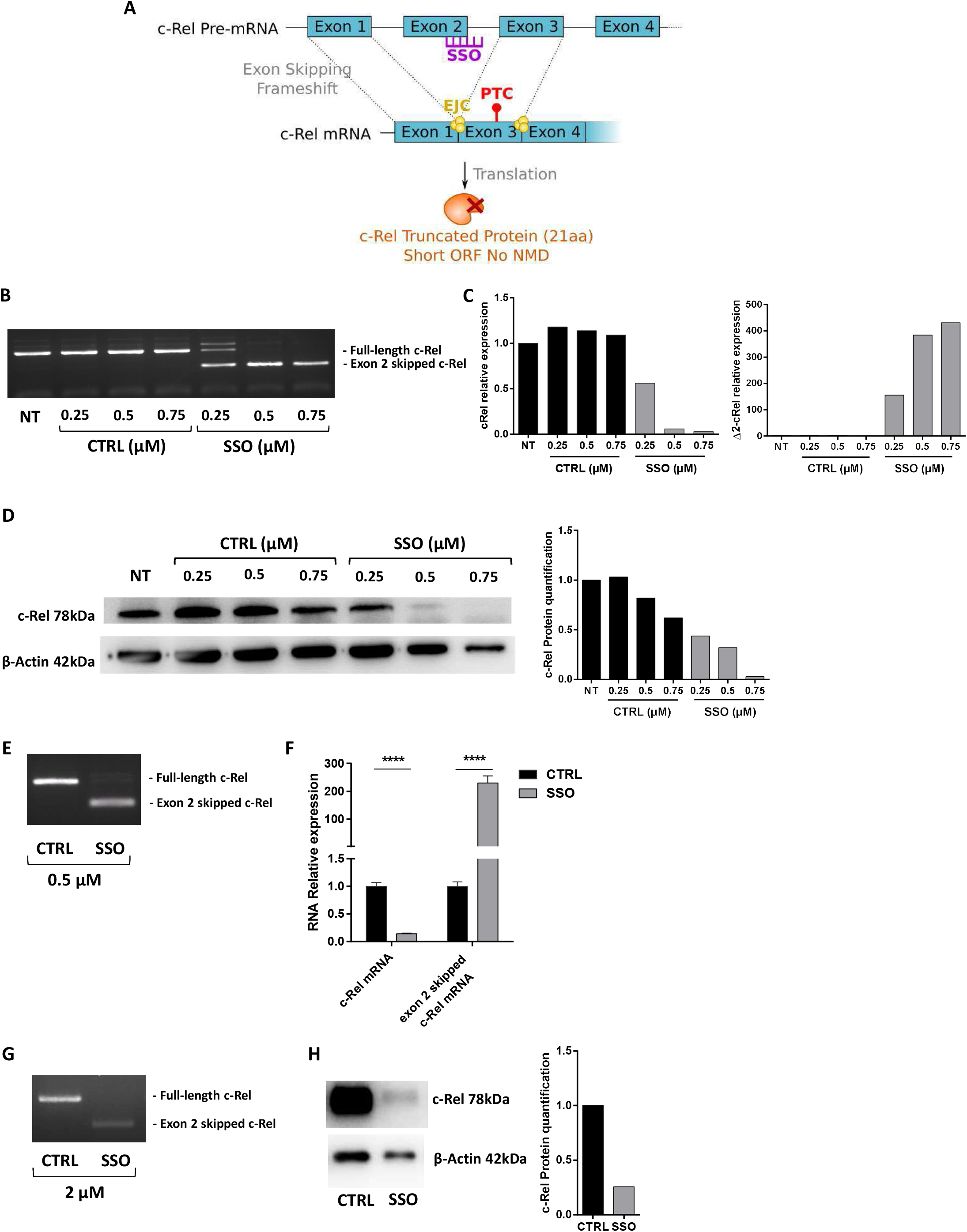
c-Rel exon 2 donor splice site targeting vivo-morpholino SSO induces efficient c-Rel knockdown in DLBCL cell lines and human plasmablasts differentiated from primary B cells. **A**, Illustration of SSO mediated knockdown of c-Rel transcripts. **B**,**C**,**D**, SUDHL4 cells were treated 48 hours with 0.25 to 0.75 µM of c-Rel exon 2 donor splice site targeting SSO (SSO) or irrelevant control ASO (CTRL). **B**, Exon 2 skipping was assessed by RT-PCR **C**, Knockdown of full length c-Rel mRNAs (left) and levels of alternative c-Rel mRNAs lacking exon 2 were measured by qRT-PCR **D**, Knockdown of c-Rel protein was verified by western blot. **E**,**F**, SUDHL4 cells were treated 48 hours with 0.5µM c-Rel exon 2 donor splice site targeting SSO (SSO) or irrelevant control ASO (CTRL) (n=4). **E**, Exon 2 skipping was assessed by RT-PCR **F**, qRT-PCRs were performed as in C. **G**,**H**, Human plasmablasts differentiated from primary B cells were treated for 72 hours at day 1 following stimulation with 2 µM SSO or CTRL. **G**, Exon 2 skipping was assessed by RT-PCR **H**, Knockdown of c-Rel protein was verified by western blot. **** p < 0.0001

### Treatment with c-Rel exon 2 donor splice site targeting SSO alters Nf-kB signalling and induces apoptosis of DLBCL cells

SUDHL4 cells treated with 0.5µM vivo-morpholino SSO were analysed by RNA sequencing to verify the absence of any other in-frame alternative transcripts. Only one out-of-frame junction resulting from intron retention downstream from exon 2 was detected in low amounts in SSO conditions (SSO) but was also present in the irrelevant control condition (CTRL) (Figure 4A). Simulating translation of this intron retention alternative transcript revealed the presence of a premature stop codon at position 319, making it a good substrate for NMD degradation and unable to restore partial c-Rel activity. Consistent with the increase of exon2-skipping events (Figure 4A), exon 2 reads including exon1-exon2 and exon2-exon3 junctions decreased while exon1-exon3 junctions appeared when cells were treated with SSO compared to the irrelevant control (CTRL).

**FIG 4.**
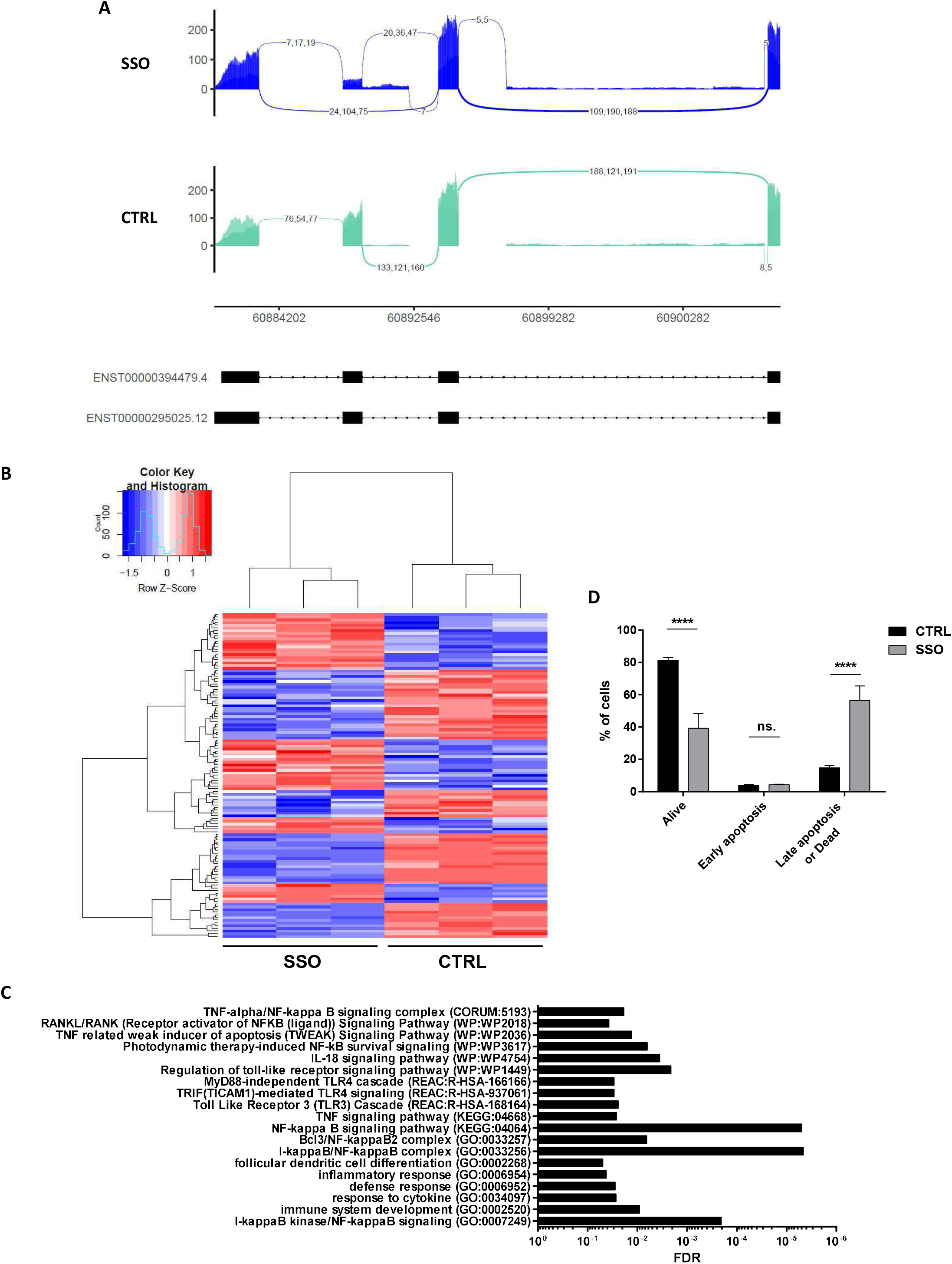
RNAseq analysis of c-Rel exon 2 donor splice site targeting SSO on splicing and biologic effects on DLBCL cells. SUDHL4 cells (n=3) were treated 48 hours with 0.5µM c-Rel exon 2 donor splice site targeting SSO (SSO) or irrelevant control ASO (CTRL) (n=3).**A**, Splice junction read analyses of exons 1 to 4 of the c-Rel gene aligned with the two known transcript isoforms. **B**, Heatmap of 121 differentially expressed genes by DEseq2 RNA sequencing analysis. **C**, Significant Gene Ontology, KEGG, Reactome, WikiPathways and CORUM pathways performed on differentially expressed gene lists. **D**, Apoptosis was analysed by flow cytometry with annexin V and 7-AAD staining. **** p < 0.0001

RNA sequencing also revealed that 122 genes were differentially and significantly expressed in DLBCL cells treated with c-Rel SSO compared to control conditions (Figure 4B). Further pathway and gene enrichment analyses revealed that the main differentially expressed pathways were related to the NF-κB signalling pathway and immune response to pathogens (Figure 4C). Among those genes, BCL3 was reduced by almost 2 fold, RELB by 2 fold, NF-κB Inhibitor alpha (IκBα), delta (IκBδ/p100) and zeta (IκBζ) by 1.5, 1.5 and 3 fold respectively. NF-κB2 was decreased by 1.5 fold, TNFAIP3/A20, NOTCH2, CD40 and CD83 by more than 1.5 fold and BIRC3 by more than 2.5 fold while PTPN6, VNN1 and FOS increased by 1.5 fold and CNTF increased by 2 fold (Table S1). Interestingly, these major transcriptomic modifications, including a deregulated TWEAK (TNF related weak inducer of apoptosis) signaling pathway, were associated with a strong increase in cell death and late apoptosis (Figure 4D).

### RelA exon 5 acceptor splice site targeting SSO efficiently knocksdown RelA protein in B cells and plasmablasts

Another SSO targeting RelA at the exon 5 acceptor splice site was designed with the help of “Exon skipper pipeline” in order to reduce RelA expression. In that case, alternative mRNAs lacking exon 5 are predicted to be NMD targets, with a PTC located at position 460 eliciting interactions with downstream exon junction complex (EJC) components (Figure 5A). OCILY10 (DLBCL) cells was used to evaluate the efficacy of passive administration of vivo-morpholino SSO to inhibit RelA expression. We observed efficient dose-dependent knockdown of RelA expression with increasing doses of vivo-morpholino SSO (SSO) from 1.5 to 2µM compared to control conditions (Figure 5B,C). As expected, full-length RelA mRNAs drastically decreased after SSO treatment, and PTC-containing alternative mRNAs were not detected and most likely eliminated by NMD (Figure 5B). At the protein level, treatment with 2µM SSO during 48 hours resulted in a nearly complete absence of RELA proteins (Figure 5C). In addition, RelA SSO was tested at 2µM in a plasma cell differentiation model described by Le Gallou et al (37,38) and compared to irrelevant control ASO (Figure 5D,E). Again, treatment with RelA SSO diminished the amounts of full-length RelA mRNAs (Figure 5D) and proteins (Figure 5E) in plasma cells expressing RelA during their differentiation program.

**FIG 5.**
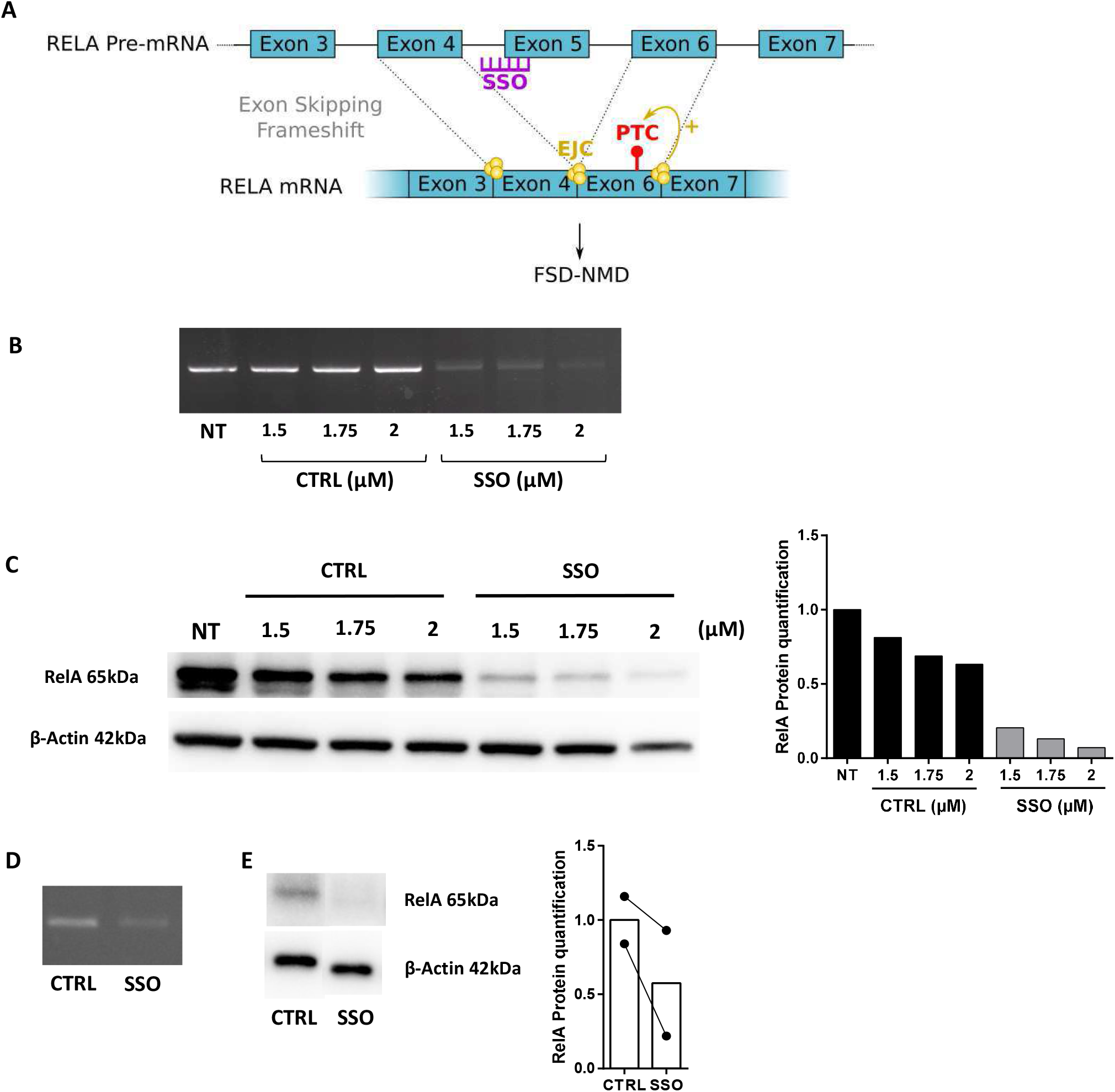
RelA exon 5 acceptor splice site targeting vivo-morpholino SSO induces protein knockdown of RelA in DLBCL cells and human plasma cells differentiated from primary B cells. **A**, Illustration of SSO mediated knockdown of RelA transcripts. **B**,**C** OCILY10 cells were treated 48 hours with 1.5 to 2 µM RelA exon 5 acceptor splice site targeting SSO (SSO) or irrelevant control ASO (CTRL). **B**, Diminution of RelA transcripts was assessed by RT-PCR. **C**, Knockdown of RelA protein was verified by western blot. **D**,**E** Human plasma cells differentiated from primary B cells were treated for 72 hours at day 4 following differentiation with 2 µM RelA exon 5 acceptor splice site targeting SSO (SSO) or irrelevant control ASO (CTRL). **D**, Diminution of RelA transcripts was assessed by RT-PCR. **E**, Knockdown of RelA protein was verified by western blot.

## DISCUSSION

An efficient transient knockdown of two differentially expressed genes during plasma cell differentiation, c-Rel and RelA, was successfully achieved with an SSO-mediated exon skipping strategy and could constitute a novel alternative to RNAi technology with known transfection efficiency limitations in primary B cells and plasma cells (10). In addition, the use of SSO could reduce off-target effects observed with RNAi technology or even RNAse-H1 dependant ASO technologies, as they require direct mRNA degradation actors : RISC/Ago2 complex and RNAse-H1 respectively. Indeed, it has been shown that as little as a 7 to 11 nucleotide homology between a siRNA and a mRNA was sufficient to recruit the RISC/Ago2 complex and induce RNA degradation, particularly in 5’ untranslated regions (39,40). This aspect remains a major issue for the development of RNAi-based therapies despite many findings in the field to reduce off-target effects using chemical modifications and optimization of RNAi structure (41). Gapmer ASOs can also recruit RNAse-H1 with unintended target binding and provoke off-target RNA degradation (42). By contrast, SSOs do not directly recruit RNA degradation machinery and therefore greatly minimize off-target effects. SSOs are also capable of weak binding to unintended sequences but the absence of RNA degradation limits the impact on off-target gene expression. However, SSOs are not completely devoid of off-target effects as splicing or RNA binding protein attachment on unintended transcripts can still be disturbed with an SSO (43).

SSO mediated c-Rel inhibition also showed interesting results in a DLBCL cell line, as it was able to reduce NF-κB signalling and cell viability. Interestingly, BCL3 expression which was described to increase cell proliferation by inducing cyclin D1 transcription (44,45) was reduced upon SSO treatment in SUDHL4 cell line. BCL3 is also known to function as an NF-κB inhibitor in the same manner as IκBζ, a Bcl-3 homolog which was also reduced after SSO treatment. It has been shown to potentiate NF-κB pathway activity by binding to p50 homodimers where they may function as co-activators (46–48). Indeed, Bcl-3 overexpression is found in chronic lymphocytic leukemia where it is frequently rearranged within the IGH locus (49). Additionally, recent findings suggest IκBζ could be a key actor in the development of psoriasis through activation of Th17 immune response (50,51) and could drive the pathogenesis of many haematological cancers (52,53). However, c-Rel is not known to have a direct effect on Bcl-3 and IκBζ expression, the latter preferentially interacting with p50 and p52 (54). Nonetheless, the *NFKB2* gene encoding p52 which is also reduced upon c-Rel targeting SSO treatment, could have influenced Bcl-3 expression as it is tightly regulated by this gene. According to our results, c-Rel was already found to be a direct regulator of NF-κB2 expression in transitional B-cells (55) and in this way could have another indirect effect on RelB expression since NF-κB2 processed protein p52 mainly associates with RelB in the alternative NF-κB pathway (56). Thus, c-Rel inhibition had a broad effect on NF-κB signalling, explaining the decreased viability found upon SSO treatment in DLBCL cells whose tumorigenicity relies on this pathway (57,58). These results still need to be confirmed in other DLBCL cell lines and patient cells to verify c-Rel knockdown benefits in this pathology but could constitute an interesting lead for the development of new innovative therapies.

Indeed, targeting the NF-κB pathway could be an effective strategy for the treatment of B-cell malignancies (59) and all NF-κB subunits (RelA, c-Rel, RelB, p52 and p50 as well as other NF-κB pathway components such as IKKs or NEMO) could be targeted by our SSO induced gene knockdown therapy. Some existing drugs have in fact been found to be active NF-κB inhibitors such as bortezomib which blocks IκB degradation (60) but no specific NF-κB inhibitor has currently shown promising results *in vivo* and furthermore they display a high risk of toxicity (61,62).

To conclude, our results show that efficient transient gene knockdown can be achieved by SSO mediated exon skipping with passive administration of vivo-morpholino oligos in primary B cells and plasma cells. Despite some limited exceptions identified with Exon skipper pipeline in mice and humans, this strategy can be implemented for most eukaryotic genes with the added advantage of a transfection-free system in primary cell cultures. Additionally, splicing modulation has the advantage of a low risk of off-target effects, as it does not rely on recruitment of direct mRNA degradation actors.

This tool could be an efficient complement to DNA modification tools such as CRISPR-Cas9 to study normal human B cells and uncover new plasma cell differentiation key mechanisms.

## Supporting information

Supplementary datas

## AKNOWLOGEMENTS

We thank BISCEm plateform (US 42 INSERM / UMS 2015 CNRS - Limoges University, Limoges, France) for the technical help in bioinformatics and transcript analysis. We thank also Claire Carrion (CNRS UMR 7276 - INSERM U 1262 - Limoges University, Limoges, France) for her help with flow cytometry experiments.

