## Supplementary datas for "Splice switching oligonucleotide mediated gene knockdown in B cells and plasma cells"

**TABLE S1. Differentially expressed gene list analysed by DESeq2 on RNA sequencing results.**

**FIG S1. Transfection of morpholino SSO targeting c-Rel exon 2 donor splice induces protein knockdown of c-Rel at high concentrations.** SUDHL4 cells were transfected with 10 to 30 µM c-Rel exon 2 donor splice site targeting morpholino SSO (SSO) or irrelevant control ASO (CTRL) and collected at 48 hours. **A,** Exon 2 skipping was assessed by RT-PCR **B**, Knockdown of c-Rel protein was verified by western blot.

**FIG S2. c-Rel exon 2 donor splice site targeting morpholino SSO transfection results in protein knockdown of c-Rel in human plasma cells.** Human plasma cells differentiated from primary B cells were treated for 72 hours at day 4 following differentiation with 2 µM c-Rel exon 2 donor splice site targeting SSO (SSO) or irrelevant control ASO (CTRL). **A,** Exon 2 skipping was assessed by RT-PCR **B,** Knockdown of c-Rel protein was verified by western blot.

#### Figure S1

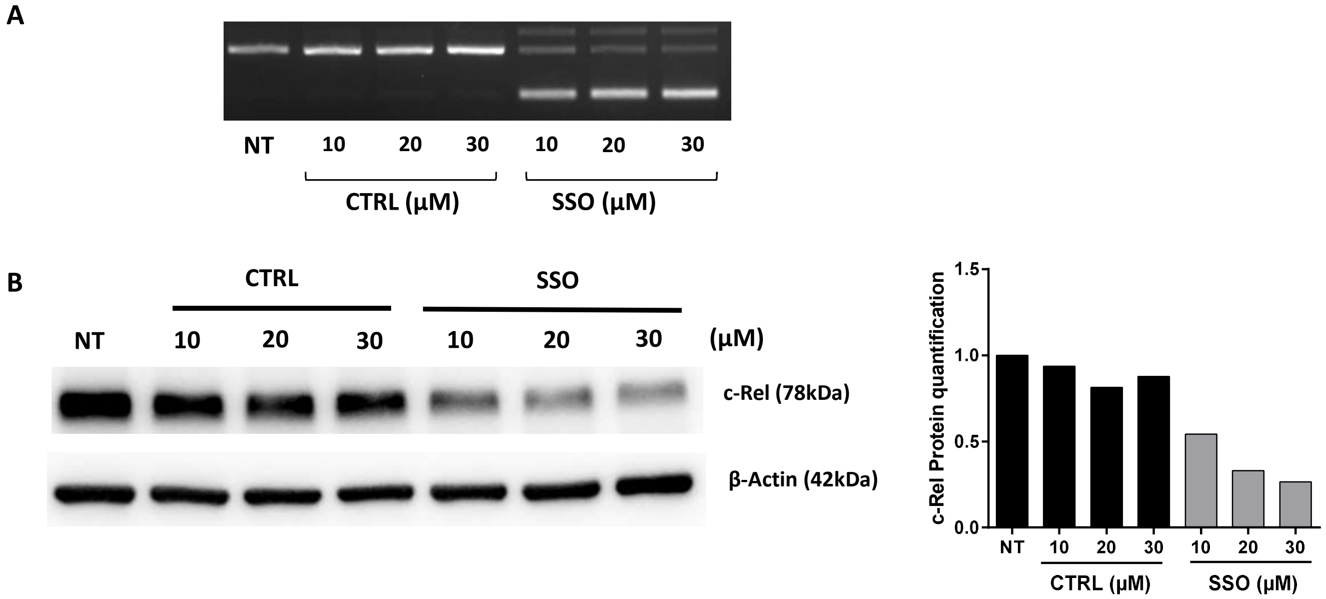


#### Figure S2

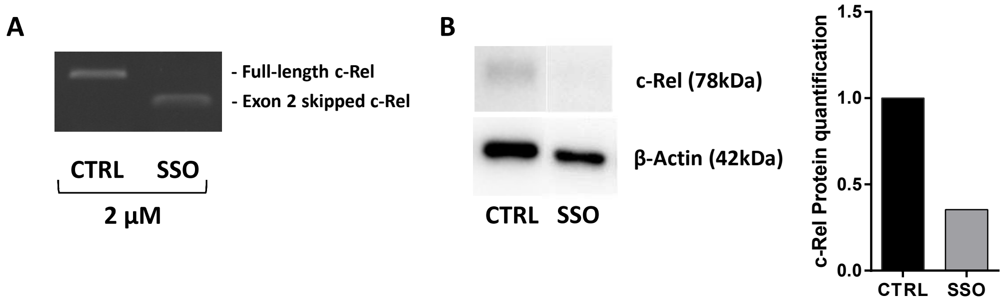

### Table S1

| Geneid | Fold change | Adjusted p value |
|---|---|---|
| C1orf132 | 5,022911295 | 7,1537E-123 |
| DOCK8 | 0,470644227 | 2,54258E-66 |
| BIRC3 | 0,37996476 | 1,53621E-56 |
| COG4 | 0,46381148 | 1,35949E-42 |
| VPS11 | 0,422596219 | 3,2518E-39 |
| RELB | 0,507445202 | 6,14813E-32 |
| POT1 | 0,521819259 | 7,0974E-31 |
| MTR | 0,50943553 | 9,72105E-31 |
| RTN3 | 0,523806126 | 6,88641E-29 |
| ECI2 | 1,918821152 | 3,65586E-27 |
| ATAD2 | 0,603364575 | 1,48395E-26 |
| CTPS2 | 0,547127552 | 1,2511E-24 |
| ZMYM4 | 0,624784662 | 4,48979E-24 |
| ADAR | 0,644808804 | 3,88417E-23 |
| KIAA0040 | 0,596174288 | 2,47178E-22 |
| MIER1 | 1,574315593 | 3,08884E-21 |
| CD83 | 0,593038068 | 5,11312E-21 |
| FBXO3 | 0,537701066 | 5,27494E-20 |
| TRAF4 | 0,661348849 | 1,2192E-19 |
| POLA2 | 0,626140895 | 1,20494E-18 |
| MIR146A | 0,353977708 | 3,30288E-18 |
| LTB | 0,378468803 | 5,71892E-18 |
| HMGCS1 | 0,617564164 | 2,7258E-17 |
| TMEM237 | 0,293582018 | 7,69124E-17 |
| ANAPC16 | 1,545896055 | 1,40711E-16 |
| DDHD2 | 0,575888194 | 3,43356E-16 |
| PEX3 | 0,616478297 | 6,27134E-15 |
| MTRR | 0,664136585 | 3,38583E-13 |
| EFCAB12 | 3,324981282 | 5,34512E-13 |
| CYB561A3 | 0,666362231 | 2,93087E-12 |
| NDUFB1 | 0,637459879 | 3,13359E-12 |
| CD40 | 0,628916087 | 1,43717E-11 |
| NBAS | 0,652615378 | 2,3053E-11 |
| MAP9 | 0,631862638 | 6,47864E-11 |
| FCRL3 | 0,610336216 | 7,98283E-11 |
| NDUFV2 | 2,884664087 | 1,03942E-10 |
| MAPKAPK3 | 0,645072101 | 1,56824E-10 |
| NFKB2 | 0,664213073 | 2,63961E-09 |
| MT-TG | 7,424980351 | 2,38681E-08 |
| WNT10A | 0,508100298 | 3,1909E-08 |
| PARVG | 1,519739422 | 1,10035E-07 |
| BCL3 | 0,573871398 | 1,354E-07 |
| NFKBIA | 0,651847682 | 1,47727E-07 |
| FAM43A | 1,495399494 | 1,61102E-07 |
| LRRN3 | 0,546433797 | 1,62289E-07 |
| NAALADL2-AS2 | 0,660553366 | 1,71901E-07 |
| CTD-2017D11.1 | 1,704172583 | 2,2197E-07 |
| TTC36 | 40,33930008 | 4,0752E-07 |
| TMEM25 | 2,594036474 | 5,68483E-07 |
| BCL2A1 | 0,551211878 | 6,8468E-07 |
| COLCA1 | 2,115020101 | 7,52887E-07 |
| GPR135 | 1,644324424 | 2,03065E-06 |
| C1orf186 | 0,667486047 | 2,48889E-06 |
| TINCR | 4,036950657 | 2,75898E-06 |
| COL24A1 | 1,739099864 | 5,93893E-06 |
| RP11-631N16.2 | 1,790169947 | 0,000031101 |
| NFKBID | 0,647982968 | 0,000034982 |
| GTF2IRD2P1 | 0,622022632 | 4,35506E-05 |
| CCDC141 | 0,594419028 | 0,000121908 |
| RP11-1072C15.4 | 2,838049507 | 0,000130096 |
| C18orf32 | 1,650368512 | 0,000150685 |
| TEKT4P2 | 1,774722385 | 0,000173249 |
| C1orf228 | 1,636140247 | 0,000223959 |
| AC004540.4 | 0,240993734 | 0,000288077 |
| CAMKV | 0,624740769 | 0,000380819 |
| GLRA3 | 1,605672679 | 0,000622837 |
| FBXO4 | 0,625584749 | 0,000655746 |
| RP11-152N13.16 | 0,487491646 | 0,00066788 |
| TMEM81 | 1,741140243 | 0,000715043 |
| SOS1-IT1 | 1,574607498 | 0,000945001 |
| IDUA | 1,62335198 | 0,001174923 |
| KLHL3 | 1,872499981 | 0,001343111 |
| OPN3 | 1,873089614 | 0,001366676 |
| ATP8B2 | 0,575910624 | 0,001398099 |
| LSMEM1 | 1,882698275 | 0,002420449 |
| CNTF | 2,123657127 | 0,002444089 |
| SH2D5 | 0,58019066 | 0,002518911 |
| SPP1 | 2,500680266 | 0,003164509 |
| VASN | 1,510811058 | 0,003286523 |
| NOTCH2 | 0,634140282 | 0,003753088 |
| MOV10 | 2,865238372 | 0,00376341 |
| KB-431C1.4 | 0,658873969 | 0,004143892 |
| ST8SIA5 | 2,184458274 | 0,005116087 |
| PNMA5 | 0,503981424 | 0,005962646 |
| KCNH8 | 1,496592874 | 0,006039196 |
| FBXO32 | 1,887811574 | 0,006106661 |
| KB-431C1.5 | 4,082214851 | 0,006278356 |
| AC079767.4 | 0,65944909 | 0,00671003 |
| RP11-381K20.2 | 0,427723137 | 0,006881883 |
| KIAA1024 | 1,705004517 | 0,006908435 |
| COLQ | 3,565504285 | 0,007040877 |
| RP11-251G23.5 | 0,619547653 | 0,007877189 |
| TNFAIP3 | 0,576843941 | 0,009515032 |
| RP1-197B17.3 | 1,680528741 | 0,0125034 |
| PARP14 | 0,339009616 | 0,014814179 |
| VNN1 | 1,734314435 | 0,017785402 |
| MGAT4A | 1,542176466 | 0,018690718 |
| RP5-1071N3.1 | 0,504132238 | 0,018812245 |
| ARHGAP24 | 0,648495529 | 0,020392585 |
| C8orf58 | 1,502131263 | 0,022456807 |
| PPP4R4 | 0,398715297 | 0,025787332 |
| ZCWPW1 | 0,623449764 | 0,02629655 |
| FOS | 1,679558968 | 0,026874758 |
| PTPN6 | 1,521024774 | 0,028168663 |
| FCRL5 | 0,482135405 | 0,029102959 |
| OXTR | 1,83248997 | 0,030425779 |
| NFKBIZ | 0,335128401 | 0,031063947 |
| MAFIP | 2,270467743 | 0,031533761 |
| SYT16 | 1,628873488 | 0,032036446 |
| OR5H6 | 0,424761529 | 0,033777723 |
| RP11-91P24.6 | 3,464564408 | 0,034763268 |
| ASXL3 | 0,510497905 | 0,034763268 |
| AVIL | 1,894696694 | 0,035959636 |
| CLEC4A | 1,727715061 | 0,036255539 |
| IFIH1 | 0,33677313 | 0,038467647 |
| MTRNR2L12 | 1,857506939 | 0,04043463 |
| TNR | 2,827475286 | 0,042379549 |
| C9orf57 | 1,655719698 | 0,042741232 |
| RP11-768B22.2 | 0,595352947 | 0,043290168 |
| ULK4 | 0,637989413 | 0,043593424 |
| TLN2 | 0,640763271 | 0,047640733 |
